# Antibody-Based Targeting of the SPP1-CD44 Axis in Pediatric High-Grade Glioma through Single-Cell and Structural Bioinformatics

**DOI:** 10.1101/2025.05.01.651763

**Authors:** Shiwani Limbu, Ambuj Kumar

## Abstract

Pediatric high-grade glioma (pHGG) is a highly aggressive brain tumor characterized by transcriptional plasticity and an immunosuppressive microenvironment. Single-cell RNA-seq analysis revealed diverse malignant and immune cell populations, with tumor-associated macrophages (TAMs) emerging as the primary source of SPP1 (osteopontin), a glycoprotein that suppresses T cell activation through CD44 binding. Cell-cell communication analysis identified the SPP1-CD44 axis as a dominant immunosuppressive pathway in the tumor microenvironment. Despite extensive transcription factor screening, no strong regulators of SPP1 were identified, suggesting regulation occurs via alternative mechanisms. To assess structural features of SPP1, replica exchange molecular dynamics simulations were performed, revealing that the CD44-binding domain is conformationally stable. Phosphorylation at Ser169, a conserved site, further stabilized this region, suggesting a potential mechanism for enhanced CD44 interaction. To disrupt this axis, among 2,500 variants of anti-SPP1 23C3 antibody, a lead candidate variant (23C3-v1) with improved SPP1 binding affinity and minimal sequence divergence was identified. Furthermore, humanized 23C3-v1 (Hu23C3-v1) was designed to neutralize Class I and Class II epitope hits from murine antibody derived 23C3-v1 antibody. Together, this study integrates transcriptomic and structural bioinformatics approaches to the target SPP1-CD44 axis, which can help reduce immunosuppressive characteristics of pHGG tumor microenvironment (TME).

## 1. Introduction

Pediatric high-grade glioma (pHGG) is an aggressive and highly heterogenous brain tumor, originating from the glial cells in brain [1]. Single-cell transcriptomic studies have shown that malignant pHGG cells recapitulate neural lineage programs, falling into four main cell states: astrocyte-like (AC-like), oligodendrocyte progenitor–like (OPC-like), neural progenitor–like (NPC-like), and mesenchymal-like (MES-like) [1,2]. These states correspond to earlier bulk transcriptional subtypes (classical, proneural, mesenchymal) but can coexist within one tumor [3]. For example, AC-like and MES-like populations often dominate “classical” and “mesenchymal” GBMs respectively, whereas OPC/NPC-like cells are enriched in “proneural” tumors [2]. Spatial transcriptomics confirms that these states localize to distinct niches - e.g. MES-like cells cluster around hypoxic, necrotic regions, while OPC/NPC-like cells populate infiltrative edges [3]. Importantly, GBM cells can transition between states (plasticity), which complicates therapies targeting any single subtype [2]. These transitions also create intermediate cell states adding another layer of complexity.

pHGG is characterized by its extensive immunosuppressive microenvironment [4]. Here, tumor-associated macrophages (TAMs) dominate the immune infiltrate, often outnumbering T cells and other lymphocytes [5]. These TAMs usually adopt immunosuppressive phenotypes that hinder T cell activation and cytotoxic function in pHGG [6,7]. Previous studies have also identified relationship between macrophage and immune activation in lower-grade gliomas (LGG), which then transitions to an immunosuppressive state in HGG [8]. One of the key mediator in crosstalk between macrophage and T cells is through Secreted Phosphoprotein 1 (SPP1), also known as osteopontin (OPN) [8,9]. SPP1 is a glycoprotein that binds the CD44 receptor on T cells, acting as an immune checkpoint that downgrades T cell activation [10,11] in both adult and pediatric gliomas. TAMs with high SPP1 expression is linked to impaired T cell responses and poor patient outcomes [11]. It has been shown that when T cells are stimulated in culture in the presence of recombinant SPP1, their proliferation and activation are markedly inhibited [12]. SPP1 causes a dose-dependent reduction in T cell division [12]. It also suppresses the production of key cytokines like interferon-gamma (IFN-γ) by activated T cells [12]. Notably, the presence of SPP1 leads to fewer T cells expressing early activation markers such as CD69 and IL-2 receptor α (CD25) after stimulation [12]. Lack of interferon signaling has shown to suppress CAR T cell activity by creating a SPP1 dependent immunosuppressive tumor microenvironment [13]. Adding anti-SPP1 antibody i.e. bioXcell Clone 100D3 and clone MPIIIB10 showed reversal of immunosuppression TME and enhanced CAR T cell antitumor responses [13]. This suggests that SPP1-CD44 interaction is key to modulating immunosuppressive TME, whose blockade could enhance T cell activity and improve responses to checkpoint inhibitors [14].

Beyond immunosuppression, SPP1-CD44 signaling in glioblastoma also promotes tumor aggressiveness by promoting stem cell–like traits and radiation resistance in glioma cells within the perivascular niche [15]. In addition, elevated SPP1 in the tumor microenvironment has been correlated with more extensive TAM infiltration in cancer [16], suggesting SPP1 may also act as a chemoattractant to recruit or retain macrophages. This dual role makes the SPP1-CD44 axis a compelling therapeutic target in pHGG. In this study we aim to combine single cell RNA sequencing data analysis and structural bioinformatics approaches to construct humanized antibody to target SPP1 protein.

## 2. Material and Methods

### 2.1. Data collection

Publicly available single-cell RNA-seq data for pediatric high-grade glioma were downloaded from the Single-Cell Pediatric Cancer Atlas Portal (ID: SCPCP000001) [17]. Only tumors obtained from patients ≤18 years of age were retained (13,663 glioblastoma cells and 1,061 other pHGG cells). Downstream analysis of raw count data is performed using Seurat v5.0 [18]. SPP1 homologues protein sequences were obtained using Blastp [19]. Homologous sequences were realigned for visualization using COBALT [20].

### 2.2. Single-cell preprocessing and cell type annotation

Cells with less than 200 expressed genes, or more than 5000 expressed genes, or with percentage of mitochondrial gene expressed more than 20 were filtered out. The top 2,000 variable genes per sample were identified for downstream analysis. Cell-cycle effects were calculated using Seurat CellCycleScoring function which implements scoring strategy described in Tirosh et. Al (2016) [21], and subsequently regressed out during data scaling using Seurat ScaleData function. Sample integration was performed using Harmony, based on the top 50 principal components (PCs). Clustering of integrated cells was then performed using 30 PCs. Differentially expressed genes for each cluster were obtained using Seurat FindAllMarkers function. Canonical marker genes, together with gene set enrichment analysis (GSEA) conducted using gseGO (ClusterProfiler v4.6) [22], was used to assign 11 cell types to all cells: T cells, NK cells, naïve B cells, microglia-derived TAM (MGD TAM), MGD transitioning macrophage (M1/M2), bone-marrow-derived (BMD) macrophage, MES-APC-like, MES-AC-like, MES-AC-like-cycling, OPC-like, and undetermined cells.

### 2.3. Copy Number Variation (CNV) Inference

InferCNV tool [23] was applied to estimate large-scale chromosomal copy number variations (CNVs) from single-cell transcriptomic data to distinguish malignant from non-malignant cell populations based on genomic instability patterns. NK cells, being non-malignant immune cells with stable genomes, were used as the reference population. The expression data was processed using default inferCNV parameters, with cutoff value 0.1. CNV profiles were then visualized to confirm tumor-associated genomic aberrations across identified cell types.

### 2.4. Cell-cell communication analysis

CellChat [24] was used to estimate cell-cell communication and their associated ligand-receptor pairs. Pathway-level communication was inferred by summarizing ligand-receptor interaction probabilities for each signaling pathway using computeCommunProbPathway(). Finally, the overall communication network was aggregated using aggregateNet() by counting the number of interactions and summarizing their communication probabilities.

### 2.5. Gene-regulatory network inference

Transcriptome-wide TF-gene target relationships were reconstructed using pySCENIC [25,26]. Homo sapiens 1390 curated TFs were used to create TF-gene pair adjacency matrix. hg38 Refseq_r80 SCENIC+ mc_v10_clust databases (https://resources.aertslab.org/cistarget/databases/homo_sapiens/hg38/refseq_r80/mc_v10_clust/gene_based/) [RRID:SCR_024808] is then used with pySCENIC ctx tool to compute the regulon enrichment matrix. The pySCENIC aucell tool was then used to generate a loom file containing regulon enrichment scores and Area Under the Curve (AUC) values. TF activity with AUC > 0.5 and Spearman correlation > 0.5 with SPP1 expression in MGD-TAM, MGD-macrophage, or MD-macrophage populations were considered strong candidate regulators of SPP1 in macrophage subsets.

### 2.6. Molecular modelling and Solute tempered replica exchange molecular-dynamics (st-REMD) simulation of SPP1 protein

Amino acid sequence of SPP1 protein was obtained from UniProt (P10451) [27]. Initial 3D protein structure of SPP1 was modelled using I-TASSER-MTD [28]. The full-length human SPP1 protein model was subjected to solute tempered replica exchange molecular dynamics (st-REMD) simulations at 283.15 K, 303.15 K, 333.15 K, and 353.15 K (four replicas, 200 ns each), using GROMACS 2023.2 [29,30] and the CHARMM36 force field [31]. Systems were solvated in a rectangular box with TIP3P water molecules at 10 Å marginal radius and neutralized by added potassium ions (K+) and chloride (Cl-) ions. All forcefield parameters and solvated protein-solvent systems for st-REMD simulation was generated using CHARMM-GUI webserver [32]. Initial energy minimization was performed using the steepest descent algorithm for 5,000 steps or until a convergence criterion of 1000 kJ/mol/nm was reached. To maintain structural integrity during minimization, position restraints were applied with force constants of 400 kJ/mol/nm² on backbone atoms and 40 kJ/mol/nm² on side chains. Electrostatics were treated with Particle-Mesh Ewald (PME) [33]. Van der Waals interactions were computed using a force-switch modifier between 1.0-1.2 nm. Systems were then equilibrated using a 4 fs timestep under NVT ensemble conditions for 10 nanoseconds (ns), with position restraints maintained on heavy atoms. Initial velocities were generated from a Maxwell distribution corresponding to the reference temperature. Electrostatics and van der Waals settings were identical to the minimization step, and hydrogen bonds were constrained using LINCS [34].

Four replicates of 200 ns simulation with solute temperature set at 283.15K, 303.15K, 333.15K, and 353.15K respectively were initiated with solvent temperature maintained at 303.15K across all replicates, where each replicate was allowed to switch every 5 ns. Pressure was maintained at 1 bar using isotropic C-rescale barostat [35]. Electrostatics were treated using Particle Mesh Ewald (PME) method [33]. Conformational stability was evaluated using per-residue root mean squared fluctuation (RMSF) score across all replicates. The quadratic mean of RMSF values [36] was computed across the three replicates to condense them into a single representative value, enabling Easy visual comparison between the reference and modified structure trajectories. An energy landscape heatmap was generated based on root mean square deviation (RMSD) and radius of gyration (Rg) values across all replicates. A representative protein conformation was selected from a global energy minimum on the energy landscape for downstream analyses.

### 2.7. Phosphorylation site prediction

NetPhos 3.1 [37] was used to predict phosphorylation site on SPP1 protein. Residue with score >0.8 was picked as a likely phosphorylation site for downstream analysis.

### 2.8. MD simulation of unmodified and phosphorylated SPP1

The relaxed SPP1 structure obtained from st-REMD was used as the starting point for conventional molecular dynamics (MD) simulations. Phosphorylation was introduced at residue S169 (pSer169), based on prediction results, and force field parameters for both the unmodified and pSer169 SPP1 structures were generated using the CHARMM-GUI webserver. Energy minimization and equilibration steps were performed under the same conditions as those used in the st-REMD protocol. Production MD simulations were carried out for both, the unmodified and pSer169 SPP1, at 303.15 K using the NPT ensemble. For each form, three independent 200 ns MD simulation replicates were conducted to ensure reproducibility. Quadratic mean of RMSF values across three replicates were calculated for each residue in unmodified and pSer169 SPP1 structure to identify protein domains affected by Ser 169 phosphorylation.

### 2.9. Antibody docking and binding affinity calculation

Complementarity-determining regions (CDRs) of parental anti-SPP1 murine antibody 23C3 [38] were identified using PyIgClassify2 [39]. 23C3 CDR region was docked to the relaxed unmodified SPP1-CD44 binding domain (residue 121-140) [40] with Haddock [41]. 2,500 23C3 antibody variants were generated with RosettaAntibodyDesign tool [42]. For every variant, full heavy and light chains sequence embeddings were obtained using ESM-2 8M model [43]. CDR-specific embeddings (H1-H4, L1-L4) were extracted from their corresponding chain full sequence embedding. Cosine distance between variant CDR sequence embedding and 23C3 CDR sequence embedding was calculated and combined with Rosetta dG_separated (SPP1-antibody binding free energy) to pick top antibody hits. Variants were visualized with UMAP generated using cosine distance obtained from embeddings. Variant with minimal dG_separated and minimal embedding distance from 23C3 was selected as the top hit variant. Mutations were mapped onto the protein complex using PyMOL [44].

### 2.10. Epitope prediction of 23C3 variant

To identify potential MHC Class I and Class II T cell epitopes, we used TepiTool [45] from the Immune Epitope Database (IEDB) [46]. Class I epitope prediction was performed using a predefined panel of the 27 most frequent HLA class I alleles, which includes both HLA-A and HLA-B supertypes: A01:01, A02:01, A02:03, A02:06, A03:01, A11:01, A23:01, A24:02, A26:01, A30:01, A30:02, A31:01, A32:01, A33:01, A68:01, A68:02, B07:02, B08:01, B15:01, B35:01, B40:01, B44:02, B44:03, B51:01, B53:01, B57:01, and B*58:01. NetMHCpan v4.1 [47] was applied to evaluate binding affinity of peptides to the selected alleles. Peptides were ranked based on predicted binding affinity (IC50), and those with an IC50 ≤ 150 nM were considered strong binders. Class II epitopes were predicted using 7-allele method using top 7 frequently observed HLA class II alleles (DRB1*03:01, DRB1*07:01, DRB1*15:01, DRB3*01:01, DRB3*02:02, DRB4*01:01, DRB5*01:01). Here, peptides were considered as strong binders if median percentile rank less than or equal to 10 [48]. Mean solvent accessibility surface area (SASA) of each epitope on 23C3-v1 structure was calculated using ShrakeRupley method [49] implemented in Biopython. All predicted class I and class II epitopes with residues mean solvent accessibility surface area greater than or equal to 50% was selected for downstream analysis.

### 2.11. Humanization of murine antibody 23C3 variant

To reduce immunogenicity while maintaining antigen-binding affinity, the murine antibody 23C3 variant was humanized through a structure- and epitope-guided approach. Initially, the top 10 humanized antibody sequences most similar to the 23C3 heavy and light chain were identified using BLASTp. The sequences of the 23C3 variant and the selected humanized antibodies were then realigned using COBALT to assess sequence conservation in class I and class II T cell epitope region. The resulting alignments were manually inspected, focusing on class I and class II T cell epitope regions and their overlap with CDR (complementarity-determining region) of the antibody. For any epitope region where the 23C3 variant sequence matched all 10 humanized antibodies, no changes were made. If differences were identified in non-CDR epitope regions, the corresponding humanized antibody amino acid sequence residue of the corresponding alignment position was introduced into the 23C3 variant sequence through point mutation to reduce potential immunogenicity. In cases where the epitope overlapped with a CDR, amino acid substitutions were made with the goal of minimizing T cell activation. Specifically, polar residues were replaced with nonpolar-hydrophobic residues to reduce MHC presentation likelihood while preserving SPP1-binding affinity, based on physicochemical compatibility with the target interface. Epitope prediction, using Tepitool, was applied on Humanized 23C3 variant sequence to test the loss of epitope from the corresponding regions.

### 2.12. SPP1 binding affinity calculation of murine and humanized 23C3 variant

Point mutations associated with successful humanization of 23C3 variants were introduced in its SPP1 docked protein structure using CHARMM-GUI. MD simulation parameters for both murine and humanized 23C3 variant structure were generated using CHARMM-GUI. Energy minimization, equilibration steps, and production MD run were performed under the same conditions as those used in Method 2.8. Three independent 200 ns MD simulation replicates were conducted to ensure reproducibility. Quadratic mean of RMSF values across three replicates were calculated for each residue in murine and humanized SPP1 variant. Binding affinity of murine and humanized 23C3 variant heavy and light chain with SPP1 across all three independent trajectories were calculated using gmx_MMPBSA [50,51].

## 3. Results and discussion

### 3.1. Single-cell landscape of pediatric HGG reveals a dominant SPP1-expressing TAM population

To define the cellular composition of the pediatric HGG microenvironment, unsupervised clustering and UMAP projection was applied on single cell RNA-seq data, leading to identified ten major cell populations (Fig. 1a), including malignant glial-lineage states (MES-like, MES-AC-like, MES-AC-like Cycling, MES-APC-like, OPC-like), lymphoid cells (T cells, NK cells, Naïve B), and myeloid cells segregating into two macrophage subsets (MGD Macrophage, MGD TAM) plus a Microglia-derived TAM cluster (MD Macrophage). A small “Undetermined” cluster did not express known canonical markers and may likely represents rare stromal elements.

**Fig 1.**
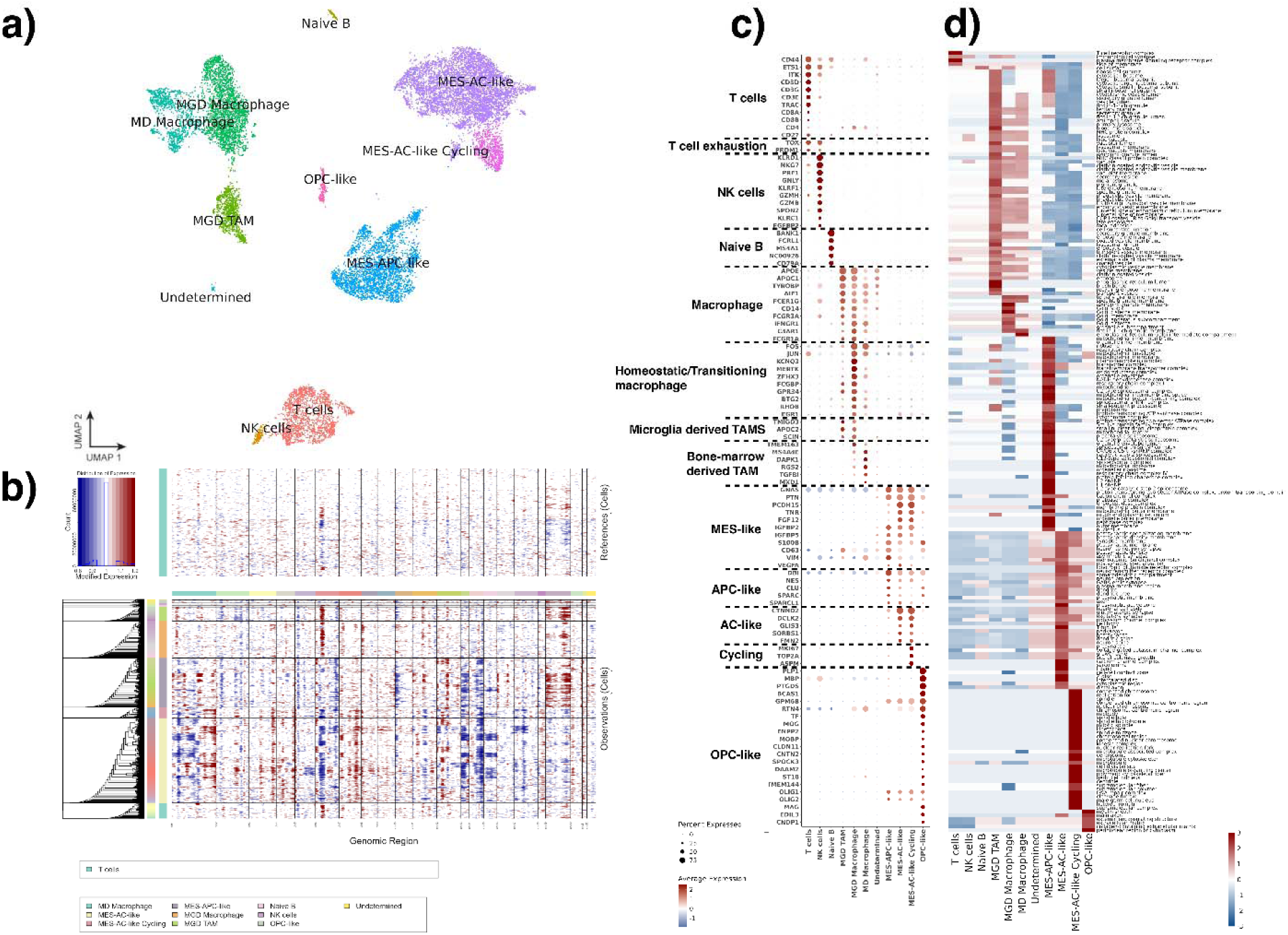
Single cell RNA-seq data analysis results. a) UMAP showing all 11 cell types identified using canonical maker gene expression profile, b) InferCNV results showing gain (red) and loss (loss) of copy number variants across all tumor cells. c) Canonical marker dotplot of commonly observed cell types in pHGG, d) GSEA results showing activated pathways across all cell types.

### 3.2. Genomic validation of malignant versus immune clusters by inferCNV

InferCNV to detect large-scale chromosomal copy-number alterations at single-cell resolution, using T cells, NK cells, and Naïve B cells as a diploid reference. The resulting CNV heatmap (Fig. 1b) revealed that all malignant glial clusters display characteristic aberration-such as chromosome 7 gain [52], as well as chromosome 1, 6, and 13 loss [53,54], while immune clusters show flat profiles consistent with a normal karyotype.

### 3.3. Gene set enrichment analysis highlights functionally distinct pathway programs in immune and tumor cell types

Single cell clusters were annotated with cell type label using their corresponding canonical gene expression markers (Fig 1c). Gene set enrichment analysis (GSEA) results show strong enrichment for the T cell receptor complex, MHC class II protein complex, and immunological synapse in T cells (Fig. 1d), consistent with an antigen-experienced, though likely restrained, T cell state within the glioma TME. Microglia-derived TAMs (MGD TAMs) showed enrichment of ribosome pathways (Fig. 1d), indicating high protein synthesis activity, which is often seen in metabolically active cells. This suggests that the MGD TAMs are actively producing proteins, possibly for cytokine secretion, antigen presentation, or immune modulation. In addition, enrichment for primary lysosome and lytic vacuole pathways suggest active engagement in phagocytosis, autophagy, or endocytosis (Fig. 1d). Furthermore, co-enrichment of MHC class II protein complex indicates that MGD TAMs are likely communicating with CD4⁺ T cells within the TME.

Homeostatic or transitioning MGD macrophages displayed an active immune modulation enrichment profile. Specifically, pathways such as tertiary granule membrane, azurophil granule membrane, and ficolin-1 rich granule membrane suggest these macrophages are trafficking immune granules and may be skewed toward an M2-like immunosuppressive phenotype. Simultaneously, the upregulation of Golgi stack, Golgi cisterna membrane, and Golgi apparatus subcompartment points to heightened protein processing activity [55], indicating functional transition in the macrophage sub population towards M2. Moreover, ficolin-enriched granule pathways imply a potential role in complement activation via the lectin pathway [56], which has dual effects in cancer, where it can either promote immune evasion through anaphylatoxins [57] or mediate tumor cell lysis [58].

MES-APC-like cells displayed a strong oxidative and translational metabolic program, suggestive of aggressive, invasive behavior (Fig 1d). Enrichment of mitochondrial inner membrane, NADH dehydrogenase complex, respiratory chain complex I, cytochrome complex, and proton-transporting ATP synthase complex indicates high oxidative phosphorylation (OXPHOS) activity [59], supporting the energy demands of invasion and proliferation. Additionally, pathways such as ribosome, mitochondrial ribosome, ribonucleoprotein complex, and rough endoplasmic reticulum indicate high translation activity. Furthermore, enrichment of spliceosomal complexes-including U2- and U12-type spliceosome, pre-catalytic spliceosome, and tri-snRNP complex-suggests that alternative splicing (AS) is actively being used in transitionary MES phenotype [60]. Ben Mrid et al., (2025) recently showed that AS is extensively rewired across glioma subtypes and associates with MES transition [61]. This raises important questions about whether splicing modulation may serve as a regulatory mechanism for glioma cell plasticity and a potential therapeutic target.

MES-AC-like cells represent a particularly intriguing transitioning phenotype. Despite lacking canonical markers of neurons or their progenitors, this cluster displays co-expression of MES-like and AC-like gene signatures while simultaneously activating multiple neuronal pathways (Fig 1d). This suggests a state of neuronal mimicry, where tumor cells may partially acquire neuron-like features, potentially facilitating immune evasion or integration into the neural niche. Another group of MES-AC like cells shows high expression of cycling cell markers (Fig 1c) as well as signatures of active mitosis (Fig 1d). Enrichment of condensed chromosomes, mitotic spindle, spindle midzone, cleavage furrow, and centriole point to cells in active cell division, most likely during S/G2/M phases of the cycle. Supporting this, activation of the kinesin complex, microtubule cytoskeleton, and microtubule organizing center indicates that these cells are undergoing chromosome segregation and cytokinesis. Furthermore, enrichment of nuclear replication fork, heterochromatin, and DNA repair complex pathways underscores ongoing DNA replication and repair, likely reflecting the heightened stress and genomic instability associated with rapid tumor cell proliferation. Collectively, gene enrichment signatures across all cancer cell type indicates that TME is actively under transition state promoting immunosuppressive phenotypes.

### 3.4. Cell communication analysis pinpoints SPP1-CD44 as a key immunosuppressive axis

To interrogate intercellular communication, we applied a Cellchat to all clusters (Fig. 2a-d). Among outgoing signals, SPP1 ranked within the top ligands secreted by MGD TAMs (Fig. 2a), while its receptor CD44 was highly expressed on T cells (Fig. 1c). In addition, Fig 2b shows that SPP1 is the top incoming signal in T cells. Dot-plot analysis of communication probabilities (Fig. 2c) confirmed that the MGD TAM - T cell axis via SPP1-CD44 is the strongest single ligand–receptor interaction in the microenvironment, exceeding even canonical cytokine pathways such as CCL3-CCR1 or IL1B-IL1R1. Network topology mapping (Fig. 2d) revealed that MGD TAMs and MGD transitioning macrophages are the dominant senders of SPP1 signals, with negligible autocrine SPP1-CD44 loops in tumor cells. Violin plots of SPP1 transcript abundance (Fig. 2e) show that MGD TAMs and MGD transitioning macrophage cells express SPP1 at levels 3-10-fold higher than other immune and cancer cells. Together, these data demonstrate that a MGD TAM and MGD transitioning macrophage cells are principal source of SPP1 in pHGG, highlighting SPP1 as a potential therapeutic target.

**Fig 2.**
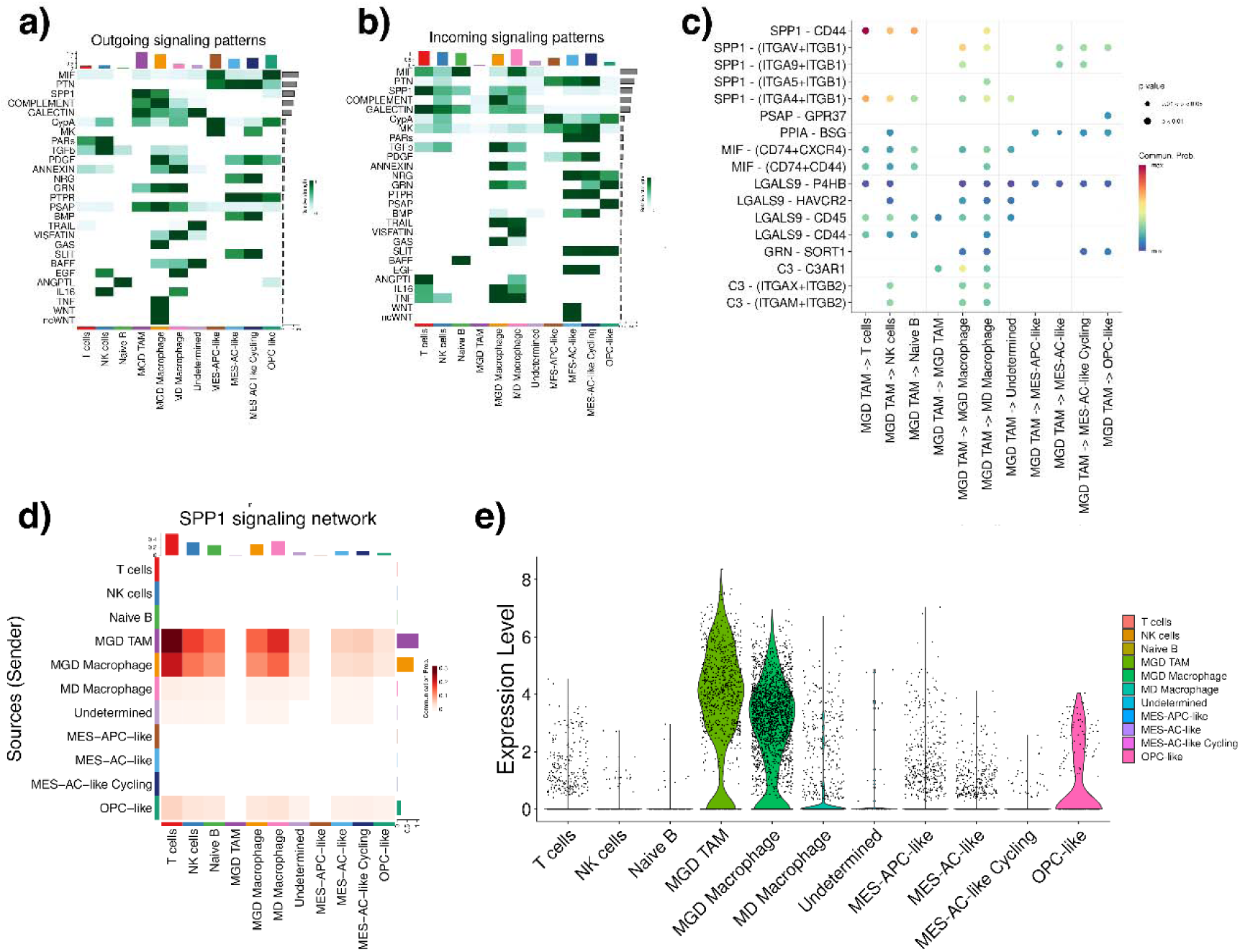
Cellchat cell-cell communication analysis results. a) Outgoing signal heatmap. The intensity of color represents higher communication probability. b) incoming signal heatmap. The intensity of color represents higher communication probability. c) All outgoing signaling network from MGD TAMs. e) Network topology heatmap of SPP1. f) Violin plots of SPP1 transcript abundance across all clusters.

### 3.5. Transcription factors regulating SPP1

pySCENIC was applied to identify transcription factors (TFs) that might drive SPP1 expression in tumor-associated macrophages (TAMs). Among six high-confidence TFs (THRB, TEAD1, RFX3, NFIB, MAFB, KLF12) predicted by pySCENIC, none exhibited strong regulon activity (AUC < 0.5), and correlation with SPP1 was uniformly weak (|r| < 0.2) in both MGD TAMs and MGD transitioning Macrophages (Fig 3a,b). MAFB stands out as highly expressed in MGD Macrophage (> 60% of cells) with high average expression, and moderately expressed in MGD TAM (Fig 3b,c,d), yet low regulon activity indicates that none of the predicted TF’s act as a dominant transcriptional driver of SPP1 in TAMs, limiting the feasibility of directly targeting these TFs to suppress SPP1 production. Therefore, alternative strategies, such as intervention at the level of upstream signaling pathways or epigenetic modulators could prove more effective.

**Fig 3.**
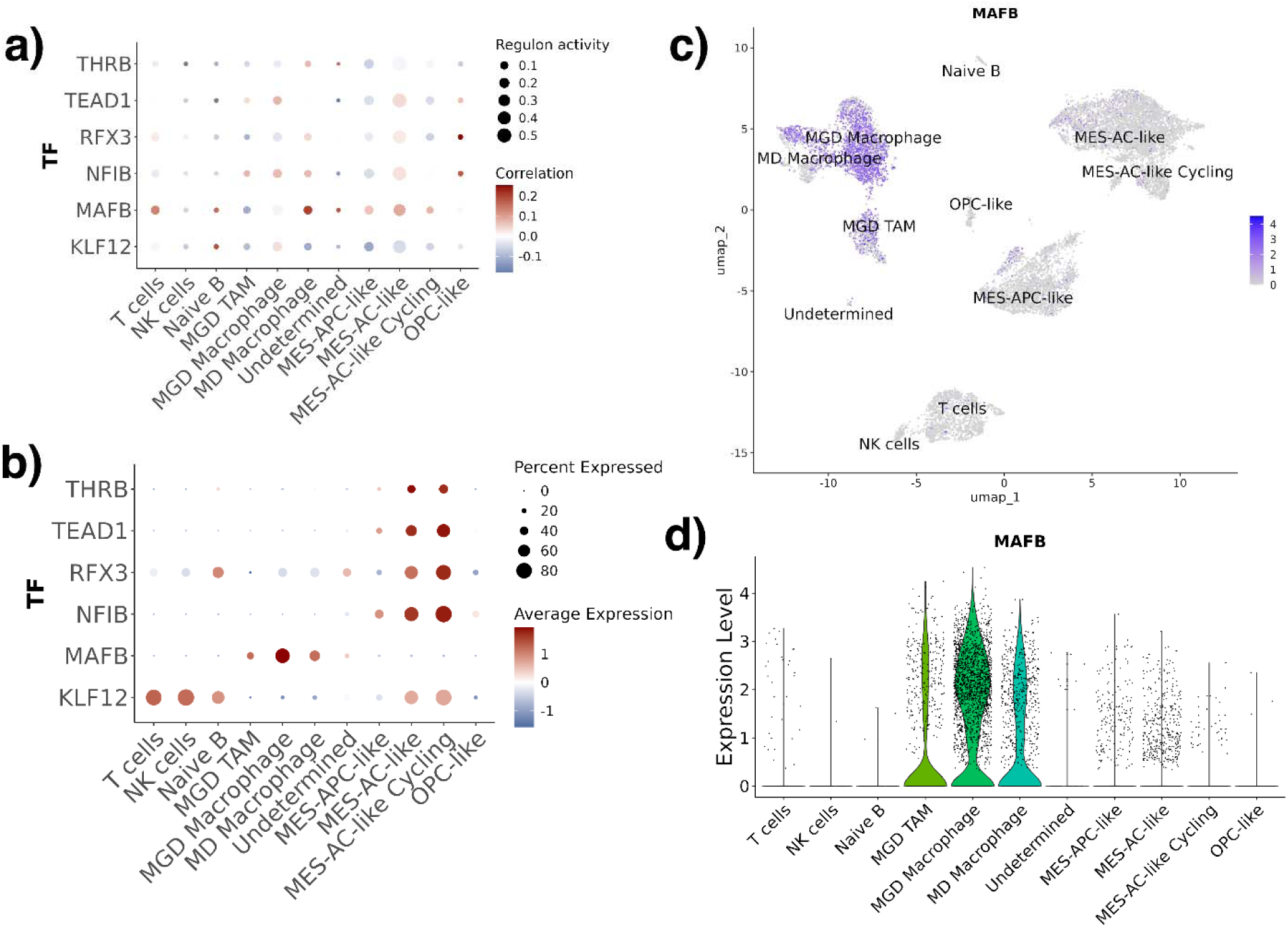
pySCENIC results highlight that no one TF dominates SPP1 gene expression regulation in MGD TAM. a) TF Regulon activity profile of SPP1 transcription factors, here size of the dot shows regulon activity and color shows gene expression correlation of the corresponding TF with SPP1 gene expression. b) TF gene expression dot plot. The size of the dot represents number of cells expressing corresponding TF in each cell type population, and the color represents average expression of the gene within the cell type population, c) MAFB gene expression feature plot showing expression of this gene across all cell types. d) MAFB violin plot showing expression of this gene across all cell types.

### 3.6. Candidate epitope regions within disordered SPP1

SPP1 is a highly disordered protein, which enables it to undergo conformational transitional associated with its diverse range of interactions in TME. Despite its intrinsic disorder, SPP1 exhibits segments of relative structural stability that has shown to serve as viable antibody epitopes [38]. Recent work showed that using an anti-SPP1 antibody (bioXcell Clone: 100D3) and (clone MPIIIB10) in combination with mIL13Rα2 CAR T cell therapy shows enhanced CAR T cell antitumor response [13]. Murine antibody 2K1, or its chimeric form C2K1, recognizes the equivalent epitope region of human SPP1 (162SVVYGLR168) [62]. Another murine anti-SPP1 antibody, 23C3, binds to residue segment 42-48 [38]. Therefore, antibody-dependent targeting of the CD44-binding domain of SPP1 holds therapeutic promise.

Due to the disordered nature of SPP1 protein, we first applied replica-exchange MD simulation at four temperatures (283 K, 303 K, 333 K and 353 K) to identify relative thermodynamic stability of different regions of the protein. Residue-wise RMSF profiles revealed extensive fluctuations across the protein, reflecting temperature-dependent modulation of its dynamic regions (Fig. 4a). Antibody binding regions 2K1, C2K1 (residue 162-168), and 23C3 (residue 43-48), consistently showed significantly lower fluctuation across replicas (Fig 4a) compared to the highly dynamic remainder of the protein. Importantly, CD44-binding domain (residue 121-140) displayed comparable stability to these known candidate epitope loops, suggesting it may be conformationally accessible for antibody binding. To capture a representative SPP1 conformation, all MD frames from all replicas were projected onto a two-dimensional free-energy surface defined by RMSD and radius of gyration (Rg) (Fig. 4b). The global free-energy minimum corresponded to a compact ensemble centered at RMSD ≈ 1.9 nm and Rg ≈ 4.09 nm. The centroid structure was extracted from this basin (Fig. 4c) for downstream analyses.

**Fig 4.**
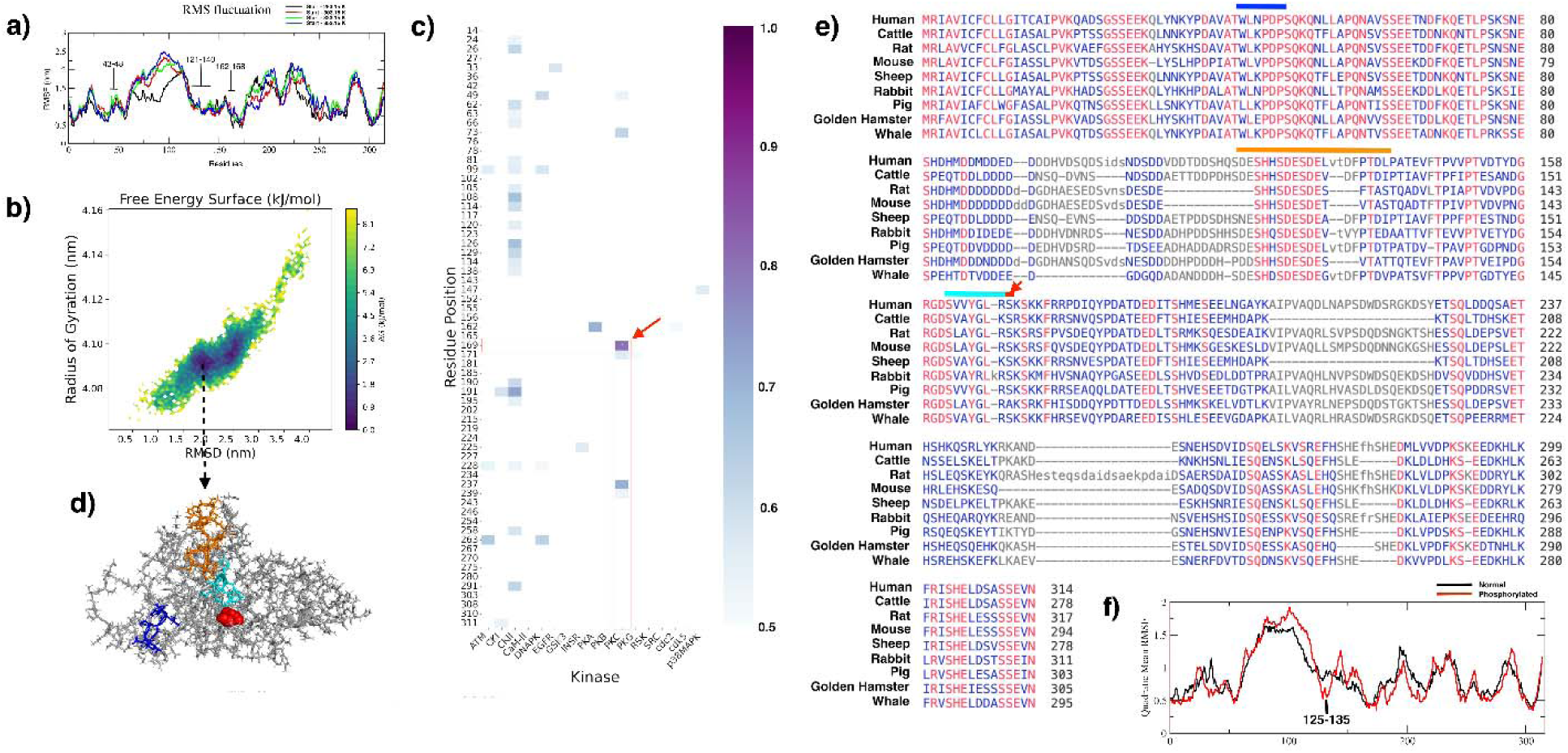
SPP1 protein structure and sequence analysis results. a) Root mean squared fluctuation of SPP1 protein residues under replica exchange molecular dynamics simulation starting at temperatures 283.15K (black), 303.15K (red), 333.15K (green), and 353.15K (blue). 8, b) Energy landscape-based conformation sampling from four replicate exchange runs. Here y-axis represents radius of gyration values across all 4 replicates and x axis represents RMSD values across all 4 replicates. c) Most stable SPP1 conformation across all 4 replica exchange trajectories. CD44 binding motif (residue 121-140) is shown in orange, 23C3 binding motif is shown in blue, and 2K1 and C2K1 binding motif is shown in cyan. and known SPP1 antibody binding interface shown in blue, and cyan, d) SPP1 protein residue phosphorylation score heatmap. Arrow highlights the top computationally predicted likely phosphorylation site 169 with score > 0.8, e) Sequence alignment of SPP1 across mammals. Orange bar highlights SPP1-CD44 binding motif, blue bar highlights 23C3 binding motif, cyan bar highlights 2K1 and C2K1 binding motif, and red arrow represent phosphorylation site 169 on SPP1 protein sequence. f) Root mean squared fluctuation quadratic mean of normal SPP1 (black) and residue 169 phosphorylated SPP1 (red) across all three independent MD simulation replicates, showing increase in stability in SPP1-CD44 binding interface upon phosphorylation.

### 3.7. Phosphorylation at Ser169 may allosterically stabilize the CD44-binding motif

Given the disorder nature of SPP1 protein, post-translational modifications could induce local folding and create stable binding surfaces. NetPhos 3.1 predicted a high-confidence Protein kinase C (PKC) phosphorylation site at Ser169 (score > 0.8) (Fig. 4d). Multi-species alignment (Fig. 4e) confirmed that Ser169, as well as the known epitope regions 43–48, 162–168, and the subset (residue 123–131) of CD44-binding domain, are highly conserved, indicating functional significance of their corresponding residues.

To evaluate the impact of residue 169 phosphorylation, three independent 200-ns MD replicates each of unmodified and pSer169 SPP1 were conducted. Quadratic mean of RMSF values of all three replicates for each residue position was calculated to explore difference between RMSF patterns in unmodified and pSer169 SPP1. Result indicates a marked reduction in fluctuation specifically in residues 125–135, a region that overlaps with the core CD44-binding domain (Fig. 4f). This allosteric dampening suggests that Ser169 phosphorylation may play a role in mediating SPP1-CD44 interaction, and thereby it can also promote a more ordered, antibody-accessible conformation of the CD44-binding loop. We plan to extend MD simulations of unmodified and pSer169 SPP1 to microseconds to verify how phosphorylation at residue 169 affects the SPP1-CDD4 binding interface at a longer time scale.

### 3.8. Antibody-optimization landscape and selection of a lead anti-SPP1 variant

Starting from the murine 23C3 monoclonal antibody, in-silico library of 2,500 CDR-mutated variants were assessed for i) predicted binding free energy to unmodified SPP1 (dG_separated) and ii) amino acid sequence/biophysical similarity to 23C3 within the eight CDRs (Table 1). Heavy- and light-chain ESM-2 embeddings, restricted to the H1-H4 and L1-L4 indices, were projected with UMAP (Fig. 5a). The resulting manifold shows two distinct sequence divergence paths with wide range of binding energy values. The color scale encodes for corresponding dG_separated values. Few variants with high relative SPP1 binding stability are shown as red dots. To integrate the two optimization criteria selected earlier, we plotted dG_separated against the cosine-esm2 embedding distance from 23C3 (Fig. 5b). The global minimum representing the most stable binder with minimal sequence embedding drift-lies near the origin of the density cloud (black arrow) (Fig. 5b). This variant combines the lowest computed binding energy with one of the smallest embedding distances (< 0.26), indicating relatively modest CDR remodeling was sufficient to achieve the desired affinity gain. Structural analysis of this top variant (23C3-v1) (Fig. 5c) reveals seven mutations in the heavy chain (S25D, T28D, N30K, I31R, N35M, T59L, T60V; red spheres) and ten in the light chain (R24A, A25C, E27D, N28D, I29V, Y30W, S31K, L33F, Q70T, Q89V; blue spheres). Because 23C3-v1 satisfies both high stability and least sequence divergence, it is prioritized as the lead candidate.

**Fig 5.**
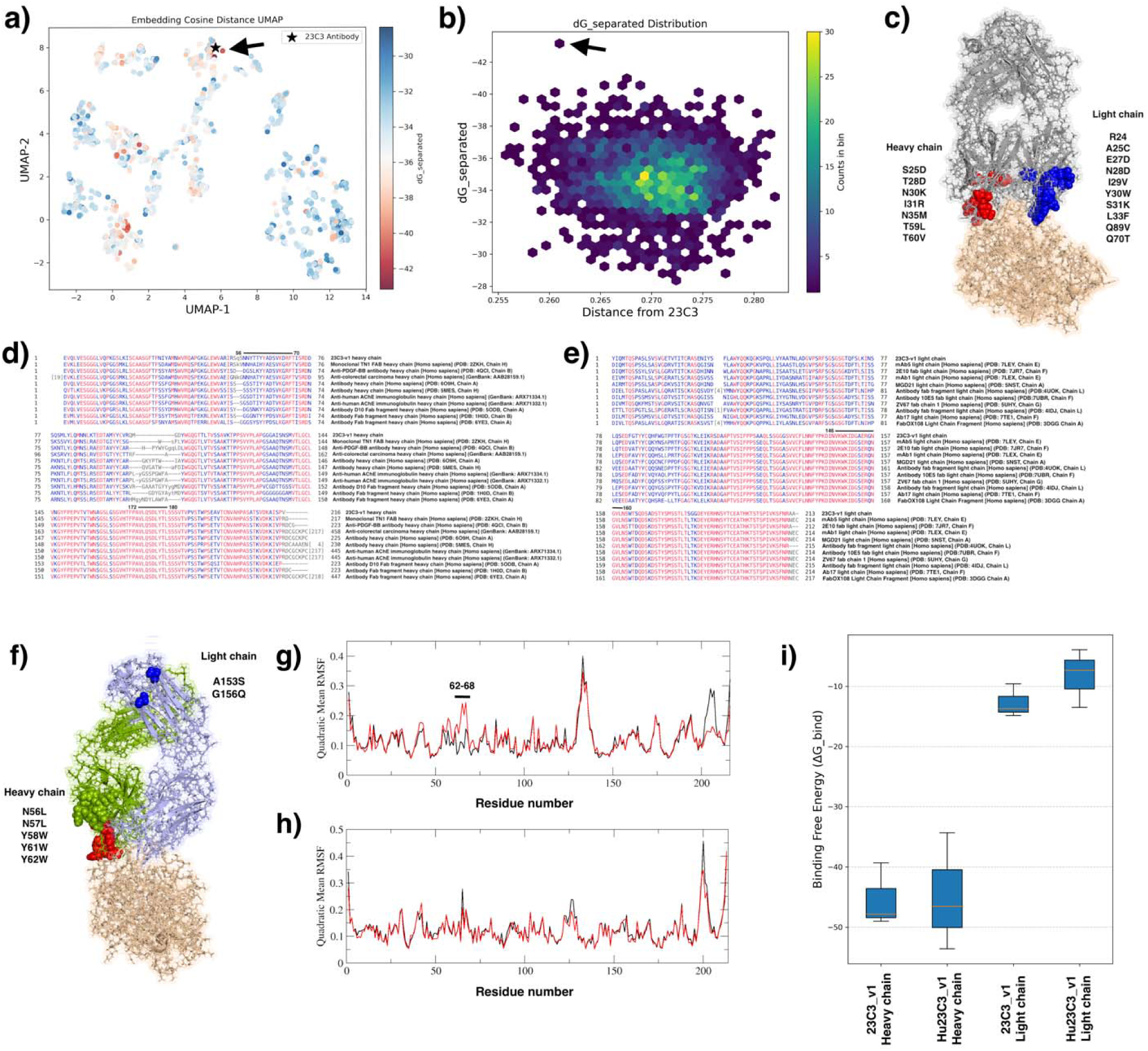
Antibody directed evolution results obtained from RosettaAntibodyDesign tool. a) UMAP showing 2500 new antibodies shown as colored dots and the parental 23C3 antibody shown as blast star. Color of the dots highlight energy required to break interaction between SPP1 and the corresponding antibody variant. UMAP is calculated using cosine distance between amino acid sequence esm2 embeddings of CDR regions of two antibody variants. b) Hex plot showing 2500 antibody variants. Here energy required to break SPP1 binding with the corresponding antibody variant is shown as x axis, and cosine distance of amino acid sequence esm2 embeddings of CDR regions of an antibody variant with parental 23C3 amino acid sequence esm2 embeddings of CDR regions. c) Most stable variant (23C3-v1) protein structure (grey) bonded to SPP1 protein structure (wheat). Blue spheres are the residue changes in 23C3-v1 as compared to 23C3 amino acid sequence. d) 23C3-v1 heavy chain amino acid sequence aligned to 10 humanized antibody heavy chain amino acid sequence. Class I and class II epitope sequence region is highlighted using black line and corresponding residue location, e) 23C3-v1 light chain amino acid sequence aligned to 10 humanized antibody light chain amino acid sequence. Class II epitope sequence region is highlighted using black line and corresponding residue location, f) Heavy chain (light green) mutations (red spheres), and light chain (cyan) mutations (blue spheres) induced in 23C3-v1 to neutralize class I and class II epitope. Heavy chain residues 62-68 are shown as green spheres, g) Three independent MD simulation replicates RMSF quadratic mean of 23C3-v1 (black) and Hu23C3-v1 (red) heavy chain, showing no significant change across SPP1 binding region of heavy chain. A sudden increase in fluctuation was observed in residue 62-68, which is distant and upward from SPP1 binding region (shown as green spheres in Fig. 5f), h) Three independent MD simulation replicates RMSF quadratic mean of 23C3-v1 (black) and humanized 23C3-v1 (Hu23C3-v1) (red) light chain, showing no significant change across SPP1 binding region of light chain, i) Box-whiskers plot of 23C3-v1 and Hu23C3-v1 heavy and light chain binding affinity with SPP1 across all three independent MD simulation replicates calculated using gmx_MMPBSA, shows no significant change in Hu23C3-v1 as compared to 23C3-v1 upon epitope neutralization.

**Table 1.** Heavy chain and light chain CDR regions of murine antibody 23C3.

| Chain | CDR region | Residue start | Residue end |
| --- | --- | --- | --- |
| Heavy chain | H1 | 23 | 35 |
| Heavy chain | H2 | 50 | 62 |
| Heavy chain | H3 | 99 | 105 |
| Heavy chain | H4 | 74 | 81 |
| Light chain | L1 | 24 | 34 |
| Light chain | L2 | 49 | 53 |
| Light chain | L3 | 89 | 97 |
| Light chain | L4 | 66 | 71 |

### 3.9. Humanization of 23C3-v1

Immunogenicity is one of the key concerns in directed evolution of a biotherapeutic protein. Sequence and structure analysis of the 23C3-v1 reveals a class I epitope (HLA class I binder) spanning residues 172–180 of the heavy chain, a class II epitope located within the engineered heavy chain CDR H2 region (residue 56-70), and a class II epitope within light chain residues 146-160 on the surface of the antibody (Table 2, Fig 5d,e). If these regions are presented on MHC II of antigen presenting cells and recognized as foreign, CD4^+^ T cells may activate, introducing helper T-cell response against 23C3-v1, and drive B-cell proliferation and differentiation leading to an anti-drug antibody (ADA) response [38]. Furthermore, presence of a class I epitope (residues 172–180) means that protein could bind an MHC I allele and be recognized by CD8^+^ T cells. CD8^+^ T cells activated by such an epitope could potentially kill other cells that present the antibody-derived peptide. Moreover, immune system clearing cells that have taken up the soluble 23C3-v1, may also lead to CD8^+^ T activation. Recently it was shown that patients treated with a CAR T, which included a murine antibody sequence, lead to CD8^+^ T cell activation, causing loss of the therapeutic population [63].

**Table 2.**
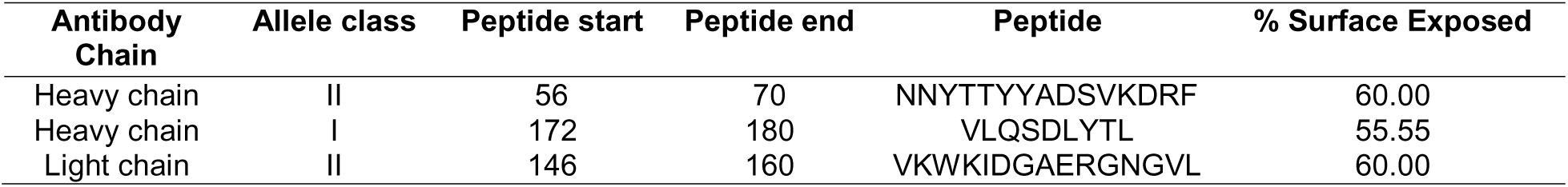
Predicted MHC Class I and II T Cell Epitope Regions in the Heavy and Light Chains of 23C3-v1.

Humanization of murine antibody is performed to reduce immunogenicity and improve safety and efficacy for therapeutic use in humans, which involves modification of its protein sequence to closely resemble that of a human antibody. Humanization of 23C3 murine antibody have been successfully implemented in Fan el., (2011), resulting in reduction of T cell response against it [64]. Therefore, directed mutagenesis was applied in the computationally predicted Class I and Class II epitope sequences of 23C3-v1 to create the humanized 23C3-v1 antibody (Hu23C3-v1). Heavy chain mutations N56L, N57L, Y58W, Y61W and Y62W (Fig 5f) helped remove residue 56-70 from Tepitool class II epitope hit. No changes in heavy chain residue 172-180 was introduced as its high conserved across all other 10 humanized antibody heavy chain (Fig 5d). Mutations A153S and G156Q were added to 23C3-v1 light chain sequence to match humanized antibody light chain sequences. Three independent MD simulation replicates of 23C3-v1 bonded to SPP1 protein, and Hu23C3-v1 bonded to SPP1 protein, were applied to evaluate change in SPP1-antibody binding affinity upon 23C3-v1 humanization. Results indicate no significant change in Hu23C3-v1 heavy and light chain RMSF as well as its binding affinity to SPP1 as compared to the RMSF and binding affinity of 23C3-v1, indicating that humanization of 23C3-v1 did not induce any deleterious impact on SPP1 binding.

Despite the encouraging in-silico affinity profile of 23C3-v1 and Hu23C3-v1, it is important to consider limitations of this study. Our 200-ns trajectories are likely insufficient to accurately capture phosphorylation induced slow allosteric rearrangements in highly disordered SPP1 protein. Moreover, all antibody modelling predictions were derived from static RosettaAntibodyDesign scoring and short-timescale MD snapshots (200 ns), which do not completely account for motions associated with conformational changes that may contribute to antibody-SPP1 binding. To address this, we will extend MD simulations to 10 micro-second timescale to confirm that PKC-mediated phosphorylation of Ser169 indeed stabilizes the CD44-binding domain, quantify how this post-translational modification modulates the free-energy landscape of the antibody–antigen complex, and evaluate impact of 23C3-v1 humanization on SPP1 binding. Furthermore, prior to experimental exploration, we aim to screen surface residues. Results obtained in this study and our follow-up analysis will provide the necessary immunogenicity, kinetic, and thermodynamic validation to advance our computationally optimized anti-SPP1 Hu23C3-v1 antibody toward experimental tests.

## Code Availability

All scripts associated with this work is available on Github (https://github.com/Sashoss/Pediatric_Immuno-Oncology)

